# Bundlemorphometry: Spatially Resolved White Matter Bundle Shape Analysis Using Nonlinear Streamline-Based Registration

**DOI:** 10.64898/2026.11.02.763786

**Authors:** Dulaj Dasanayake, Sumonta Nandy Amit, Bramsh Qamar Chandio

## Abstract

Diffusion MRI tractometry quantifies microstructural properties along white matter (WM) bundles but does not directly measure their geometry, potentially overlooking localized morphological alterations. Here, we introduce BundleMorphometry, a framework for quantifying local WM bundle shape using the analytical Jacobian of nonlinear streamline-based registration. Individual subject bundles are registered to a reference atlas bundle using BundleWarp, a streamline-based nonlinear registration method. We derive the analytical Jacobian of the resulting deformation field and compute its log-determinant (log det J) to quantify local expansion and compression at each streamline point. To enable spatially resolved group comparisons, these measurements are projected onto the two-dimensional grid, capturing variations along and across each bundle. We evaluated BundleMorphometry using two clinical datasets comprising patients with glioma and Parkinson’s disease (PD). In the glioma dataset, BundleMor-phometry revealed spatially extensive deformation patterns in lesion-affected WM bundles that were not captured by displacement magnitude alone. Additionally, in a group analysis of the PD dataset (32 patients and 32 healthy controls), we identified significant localized morphological differences in the corpus callosum forceps major, mid-body and right spinothalamic tract, with distinct patterns of expansion and compression. These findings demonstrate the potential of BundleMorphometry to advance conventional tractometry by enabling spatially resolved analysis of bundle morphology.

## 1. INTRODUCTION

Diffusion MRI (dMRI) tractography reconstructs white matter (WM) pathways as bundles of streamlines, while tractometry quantifies microstructural properties, such as fractional anisotropy (FA), mean diffusivity (MD) and related diffusion measures along these bundles [1, 2]. However, conventional tractometry primarily assesses tissue microstructure along WM pathways without directly measuring their geometry or shape. Diffusion-derived measures, such as FA, are also influenced by fiber orientation and organization within a voxel, complicating the interpretation of microstructural alterations independently of fiber geometry [3]. WM bundle geometry changes throughout development and aging [4]. In neurological disorders such as glioma, tumors may displace, infiltrate, or disrupt surrounding WM pathways [5]. Therefore, a spatially resolved measure of WM bundle morphology is needed to quantify localized geometric changes along and across WM pathways, beyond the microstructural properties captured by conventional tractometry.

Existing bundle-level shape descriptors, such as length, span, volume and irregularity [6, 7], provide global measures of WM bundle shape but do not capture spatially localized morphological variations. In structural MRI, voxel-based morphometry compares local tissue composition [8], whereas tensor-based morphometry (TBM) localizes shape differences with the Jacobian determinant of a nonlinear warp to a template, analyzed on a logarithmic scale [8, 9]. In dMRI, TBM has been extended to diffusion tensor-based registration to identify localized WM morphological differences [10], and fixel-based analysis measures morphology through the fiber cross-section [11]. These methods, however, estimate the warp on a voxel grid and not on individual bundles. The closest prior work computed the Jacobian of large deformation diffeomorphic registration of fiber-cluster centerlines using finite differences [12], but represented each tract by a single centerline streamline rather than the entire bundle.

BundleWarp [13] is a nonlinear registration method that operates directly in streamline space. It matches subject streamlines to an atlas streamlines and deforms each matched pair using a smooth Gaussian-kernel transformation, with deformation flexibility controlled by a regularization parameter *λ*. Because the deformation field for each matched streamline pair is defined in closed form, its spatial derivative can be computed analytically at every point, providing a basis for quantifying local bundle deformations. Here, we introduce BundleMorphometry, a framework that uses the analytical Jacobian of BundleWarp to quantify local expansion and compression relative to an atlas. Our contributions are (1) a point-wise measure of WM bundle deformation, *L* = log det **J**, computed at every streamline point from the closed-form Jacobian **J** of the BundleWarp deformation field; (2) spatially resolved analysis of *L* along and across WM bundles using the two-dimensional Spatial Inference for Tractometry (SPECTRA) grid [14], with cell-wise linear mixed model (LMM) and false discovery rate (FDR) correction; and (3) a demonstration using two clinical datasets: three patients from the University of California San Francisco Preoperative Diffuse Glioma MRI (UCSF-PDGM) dataset [15], and a group analysis of 64 participants from the Parkinson’s Progression Markers Initiative (PPMI) [16], including 32 with Parkinson’s disease (PD) and 32 healthy controls (CN).

## 2. METHODS

### Data and Preprocessing

**Glioma**. We evaluated the framework on three randomly selected UCSF-PDGM patients [15] to examine localized tumor-associated tract deformations: patient 1 (59F, left glioblastoma, WHO grade 4), patient 2 (31M, right IDH-mutant astrocytoma, grade 3), and patient 3 (68F, left glioblastoma, grade 4). dMRI data were acquired on a 3T GE Discovery 750 scanner with 55 diffusion directions at *b* = 2000 s/mm^2^. **PPMI**. Additionally, we tested group analysis on the 64 PPMI participants [17, 16], including 32 PD and 32 CN (age range: 39–61 years; 18 males and 14 females per group). All participants were scanned on Siemens 3T scan-ners with 2 mm isotropic voxels, 64 gradient directions at *b* = 1000 s/mm^2^ and one *b* = 0 image.

dMRI preprocessing included skull stripping, denoising, motion correction, and eddy-current correction. Fiber orientation distributions were estimated using constrained spherical deconvolution (CSD), followed by deterministic tractography. WM bundles were segmented using RecoBundles in DIPY [18, 19]. For Glioma, we selected four tracts that meet these lesions (Fig. 2), the left corticospinal tract (CST) of patient 1, the right uncinate fasciculus (UF) of patient 2, and the left arcuate fasciculus (AF) and left inferior longitudinal fasciculus (ILF) of patient 3. A streamline was considered to cross the lesion if any of its points lay inside the tumor segmentation. For PPMI, we analyzed ten bundles: the forceps major and mid-body of the corpus callosum (CC), left AF, right CST, right spinothalamic tract (STT), left optic radiation (OR), and bilateral ILF and occipitopontine tracts (OPT).

### 2.2. BundleMorphometry

#### 2.2.1. BundleWarp Registration and the Analytical Jacobian

BundleMorphometry uses BundleWarp to register each subject bundle to its corresponding atlas bundle, establishing a common spatial reference for comparisons across subjects (Fig. 1a,b). A regularization parameter of *λ* = 0.01 was used to favor close alignment with the atlas. BundleWarp first aligns the two bundles with an affine transform. It then pairs subject and atlas streamlines by linear assignment and deforms each pair with a memoryless version of coherent point drift [20], in which the *M* points **y**_*m*_ of the subject streamline act as Gaussian mixture centroids fitted to the atlas stream-line. The resulting deformation adds a displacement field **u** to every point **z**,

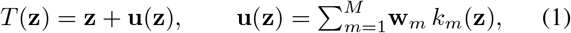

where *k*_*m*_(**z**) = exp −*∥***z** − **y**_*m*_ *∥*^2^*/*2*β*^2^ are Gaussian kernels centered on the subject points and the weights **w**_*m*_ *∈* ℝ^3^ are estimated by expectation–maximization with the penalty *λ* on the deformation. We used a kernel width of *β* = 20 mm, and **u** is the deformation field (Fig. 1c).

**Fig. 1.**
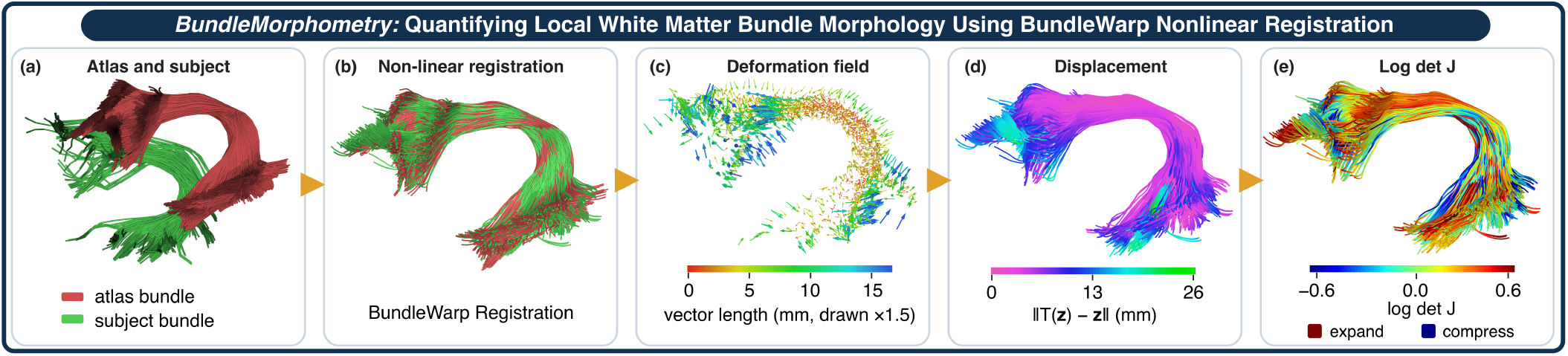
Overview of BundleMorphometry. (a) Left arcuate fasciculus (AF) atlas and subject bundles before registration. (b) Nonlinear registration using BundleWarp. (c) Estimated deformation field. (d) Displacement magnitude and (e) log det **J**, representing local expansion and compression on the registered bundle.

The Jacobian of *T* is the matrix of its first derivatives. Since each **w**_*m*_ is constant in **z**,

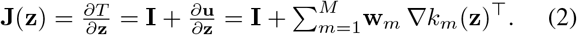

Differentiating the Gaussian kernel with the chain rule gives

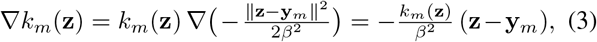

and substituting (3) into (2) yields the closed form

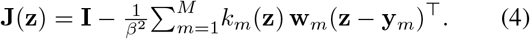

This 3 × 3 matrix depends on the kernel centers and weights that BundleWarp computes for each streamline pair, so no finite differences or voxel grid are required. BundleMorphometry evaluates (4) at every streamline point and computes the log-Jacobian determinant *L*(**z**) = log det **J**(**z**) where det **J**(**z**) *>* 0, which can be visualized on the warped bundle next to the displacement magnitude *∥***u**(**z**)*∥* (Fig. 1d,e). For small deformations, log det(**I** + **A**) ≈ tr **A**, and therefore

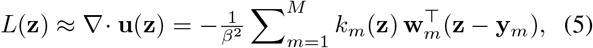

which is the divergence of the displacement field. It is positive where the field spreads points apart and negative where it draws them together.

det **J**(**z**) is the ratio of the volume of a small neighborhood of **z** after and before the deformation. Since the mapping runs from the subject to the atlas, positive values of *L* indicate that the subject bundle is locally smaller than the atlas and had to be expanded, whereas negative values indicate that it is locally larger and had to be compressed. The local volume ratio is *e*^*L*^; for example, *L* = −0.1 corresponds to a subject bundle approximately 10.5% larger in volume than the atlas at that point. A rigid translation or rotation of a bundle gives *L* = 0 everywhere, and the logarithm treats expansion and compression by the same factor symmetrically.

#### 2.2.2. Spatial Mapping and Statistics

For group-level comparisons, BundleMorphometry maps subject bundles to a common atlas space and excludes points with det **J** ≤ 0, where *L* is undefined. It uses SPECTRA [14], which enables spatially resolved analysis along and across WM bundles by dividing the atlas bundle into a two-dimensional grid along the tract (*s*) and radially from its centroid (*r*). It maps *L* onto this grid using square cells of about 1.5 mm (Fig. 3a). The value of a cell is the mean of *L* over the *n*_*i*_ valid points **z**_*j*_ of subject *i* that fall in it,

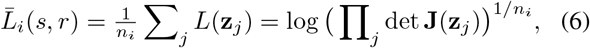

which is the logarithm of the geometric mean of det **J** in the cell. Cells with data in at least 30% of subjects and at least 10 subjects per group were analyzed with an LMM, including group, age and sex as fixed effects, and site as a random term, as in SPECTRA [14]. For each cell, we report the *p*-value of the group effect with Cohen’s *d* as the effect size, and controlled the FDR across the cells of each bundle with the Benjamini–Hochberg (BH) procedure at *q <* 0.05 [21].

## 3. RESULTS

### Computational cost

for BundleWarp and Jacobian evaluation were measured across four bundles. (Table 1).

**Table 1.**
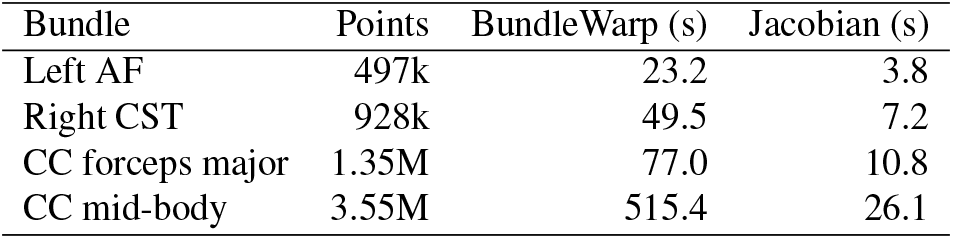
Median time per bundle over the 64 PPMI participants for BundleWarp and for evaluating the analytical Jacobian at every point in four bundles of different sizes.

### Glioma

BundleMorphometry revealed localized morphological alterations in WM bundles affected by glioma, with 65–100% of streamlines intersecting the lesions across the four analyzed tracts. In the right UF (patient 2) and left AF (patient 3), where all streamlines intersected the lesions, 80% and 90% of cells exhibited positive *L*, respectively, indicating that these bundles underwent local expansion during registration to the atlas (Fig. 2b,c). The left CST (patient 1) showed spatially varying deformation patterns, with compression in the cortical fan and brainstem and expansion in the middle segment intersecting the lesion (Fig. 2a). Similarly, the left ILF (patient 3) exhibited alternating regions of expansion and compression along the tract (Fig. 2d).

**Fig. 2.**
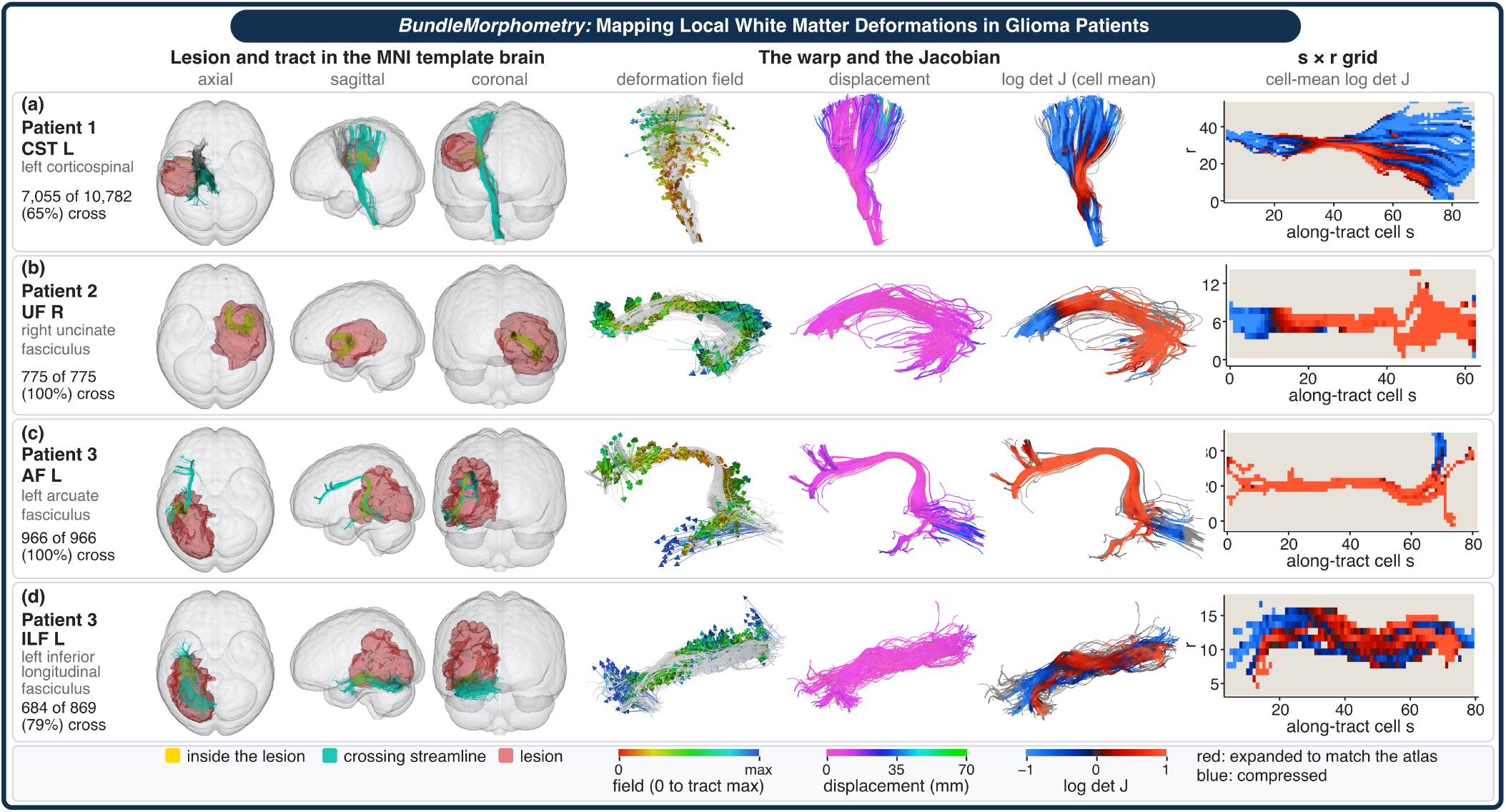
BundleMorphometry analysis of local WM deformations in glioma patients. (a) Patient 1, (b) Patient 2, (c, d) Patient 3. From left, lesion (red) and tract in the MNI template brain (crossing streamlines teal, their parts inside the lesion yellow), deformation field, displacement and cell-mean log det **J** on the tract, and the cell means on the SPECTRA *s* × *r* grid.

### PD versus Controls

Additionally, BundleMorphometry revealed spatially localized morphological differences between PD patients and CN, particularly in the CC forceps major, CC mid-body, and right STT. In the CC forceps major, 281 of 1,126 cells (25.0%) showed nominal significance (*p <* 0.05), all with lower *L* in PD, of which 20 survived BH correction within the bundle (Fig. 3b). Most of these cells were located along one occipital limb, between the splenium and the cortex. The CC mid-body showed 247 cells with nominal significance (*p <* 0.05), with 97% exhibiting higher *L* in PD. The right STT showed differences in the opposite direction to the CC forceps major, with 26 of 265 cells (9.8%) exhibiting higher *L* in PD (*p <* 0.05; Fig. 3c). The right ILF showed 61 of 511 cells (11.9%) with nominal significance, all with lower *L* in PD. Across the ten bundles, five had more than 5% of cells showing nominal significance, with at least 96% of these cells exhibiting differences in the same direction, whereas the remaining five showed fewer significant cells with mixed directions.

**Fig. 3.**
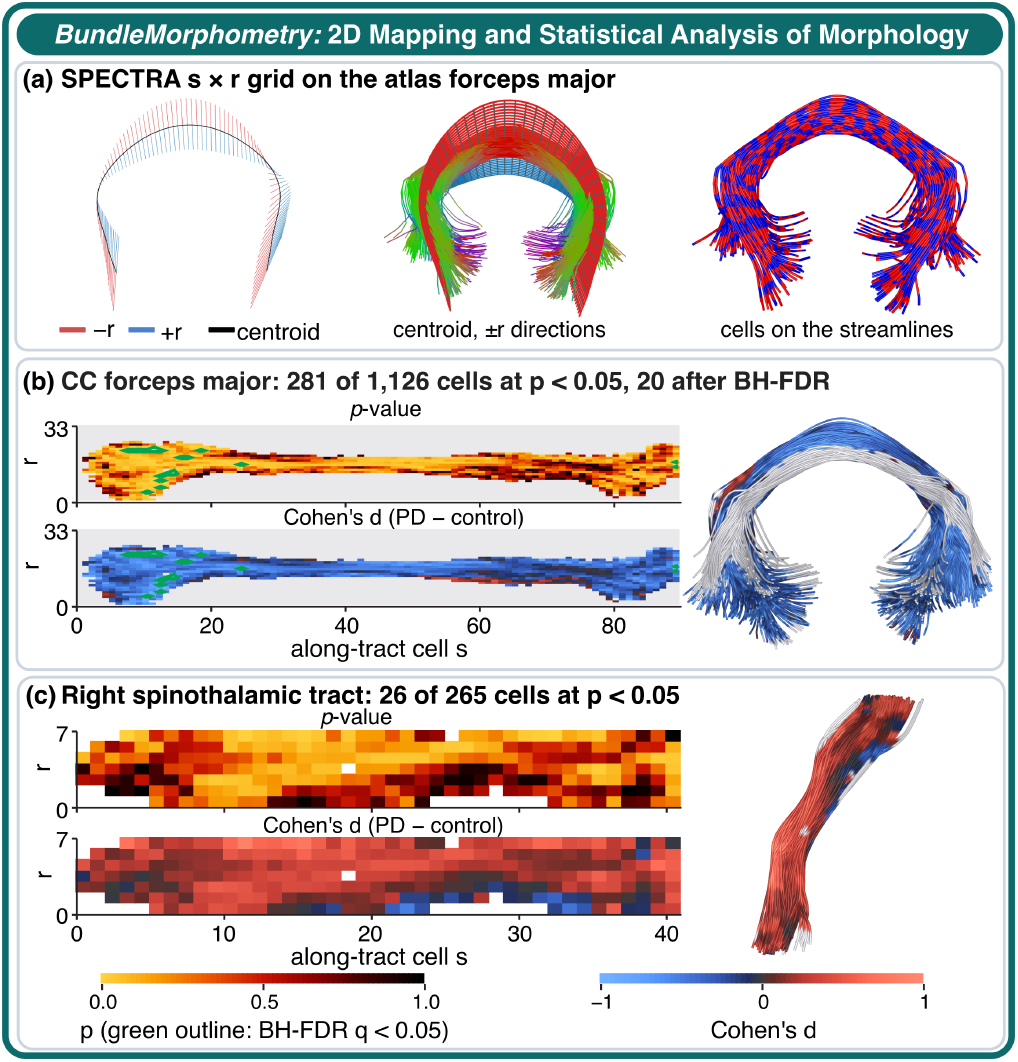
BundleMorphometry analysis of PD versus controls. (a) SPECTRA grid on the atlas CC forceps major. (b, c) LMM *p*-value of the group effect and Cohen’s *d* in each cell of the CC forceps major and the right STT, with cells that survive BH correction outlined in green, and *d* painted on the atlas bundle.

## 4. DISCUSSION AND CONCLUSION

BundleMorphometry quantifies localized WM bundle shape changes using the analytical Jacobian of BundleWarp’s non-linear streamline-based registration, capturing local geometric deformations relative to a reference atlas. The Jacobian determinant, det **J**, represents the local volume change induced by registration, while its logarithm, *L* = log det **J**, quantifies local expansion and compression relative to the reference atlas. Since all subject bundles are registered to the same atlas, differences in *L* between groups provide a measure of relative local volumetric differences in WM bundle geometry. By integrating BundleWarp with SPECTRA, BundleMorphometry enables spatially resolved analysis of *L* both along the length and radially across WM bundles. This two-dimensional representation preserves local morphological variations that may be obscured by conventional tractometry, which averages measurements across streamlines within each along-tract segment/nodes [1, 22].

In glioma patients, positive *L* in lesion-affected segments indicated local expansion toward the atlas, potentially reflecting tumor-related displacement or compression of reconstructed WM pathways, as well as changes in tractography due to infiltration or edema [5].

In the PPMI dataset, BundleMorphometry revealed localized morphological differences in the CC mid-body, CC forceps major, and right STT between PD patients and controls. The CC forceps major showed lower *L* in PD, suggesting locally larger bundle geometry, whereas the right STT showed higher *L*, suggesting the opposite pattern. These findings demonstrate the potential of BundleMorphometry to identify pathway-specific morphological differences in PD. However, larger, independent cohorts are needed to improve statistical power, detect more subtle effects, and establish the reproducibility of these findings.

Future work will evaluate BundleMorphometry in larger, independent cohorts to validate its reproducibility and generalizability across neurological disorders. We will also integrate BundleMorphometry with microstructural measures such as FA and MD to jointly assess morphological and microstructural alterations, providing more comprehensive insights into WM changes.

In conclusion, BundleMorphometry introduces a computationally efficient, spatially resolved framework for quantifying WM bundle morphology. Our findings demonstrate its ability to detect localized morphological differences in PD and lesion-associated deformations in glioma, introducing a new dimension to tractometry by enabling the study of bundle geometry beyond conventional microstructural measurements. BundleMorphometry will be available through DIPY.

## 5. COMPLIANCE WITH ETHICAL STANDARDS

This study was conducted retrospectively using open-access human subject data from PPMI and UCSF-PDGM. Ethical approval was not required as confirmed by the licenses attached with the open-access data.

## 6. ACKNOWLEDGMENTS

This work was supported by the National Institute of General Medical Sciences (NIGMS) of the National Institutes of Health (NIH) under Award Number 5U54GM104942-08.

